# Competitive fitness and homologous recombination of SARS-CoV-2 variants of concern

**DOI:** 10.1101/2023.07.26.550688

**Authors:** Qi Chen, Si Qin, Hang-Yu Zhou, Yong-Qiang Deng, Pan-Deng Shi, Hui Zhao, Xiao-Feng Li, Xing-Yao Huang, Ya-Rong Wu, Yan Guo, Guang-Qian Pei, Yun-Fei Wang, Si-Qi Sun, Zong-Min Du, Yu-Jun Cui, Hang Fan, Cheng-Feng Qin

## Abstract

SARS-CoV-2 variants continue to emerge and cocirculate in humans and wild animals. The factors driving the emergence and replacement of novel variants and recombinants remain incompletely understood. Herein, we comprehensively characterized the competitive fitness of SARS-CoV-2 wild type (WT) and three variants of concern (VOCs), Alpha, Beta and Delta, by coinfection and serial passaging assays in different susceptible cells. Deep sequencing analyses revealed cell-specific competitive fitness: the Beta variant showed enhanced replication fitness during serial passage in Caco-2 cells, whereas the WT and Alpha variant showed elevated fitness in Vero E6 cells. Interestingly, a high level of neutralizing antibody sped up competition and completely reshaped the fitness advantages of different variants. More importantly, single clone purification identified a significant proportion of homologous recombinants that emerged during the passage history, and immune pressure reduced the frequency of recombination. Interestingly, a recombination hot region located between nucleotide sites 22995 and 28866 of the viral genomes could be identified in most of the detected recombinants. Our study not only profiled the variable competitive fitness of SARS-CoV-2 under different conditions, but also provided direct experimental evidence of homologous recombination between SARS-CoV-2 viruses, as well as a model for investigating SARS-CoV-2 recombination.

**Importance:** SARS-CoV-2 variants or subvariants keep emerging and the epidemic strains keeps changing in humans and animals. The continued replacement of the epidemic strains was attributed to higher competitive fitness evolved by the newly appeared ones than the older ones, but which factors affect the final outcomes are still not entirely clear. In this study, we performed in vitro coinfection and serial passage with three VOCs and WT under different conditions. Our results showed that the competition outcomes of these viral strains varied in different cell lines or under different immune pressure, confirming the probable effects of these two factors for the competitive fitness of different SARS-CoV-2 viral strains. Meanwhile, strikingly, we found that coinfection and serial passage with different SARS-CoV-2 viral strains can mimic the recombination process of SARS-CoV-2 occurred in coinfection individual, indicating it is a novel model to investigate the SARS-CoV-2 recombination mechanism.

## Introduction

During the past 3 years of the COVID-19 pandemic, numerous SARS-CoV-2 variants have emerged from different geographic regions, and the cocirculation and displacement of different variants have been well documented worldwide. The World Health Organization (WHO) has classified and characterized a panel of specific Variants of Concern (VOCs). Briefly, the Alpha, Beta or Gamma VOCs replaced the wild-type (WT) SARS-CoV-2 and other prior strains depending on the geographic region, and these VOCs were all further replaced by the subsequent Delta variant(1-4). These VOCs have now been replaced worldwide by the most recent Omicron variant and its sublineages(5, 6). The constant replacement of SARS-CoV-2 prevalent variants is thought to be determined by multiple factors, including viral fitness, host immunity and public health policy(7); however, the intrinsic mechanism remains largely unknown.

Viral fitness is a determinant of adaptive evolution and variant emergence of several pandemic viruses, such as influenza A virus (IAV) and West Nile virus (WNV) (8-10). After the appearance of SARS-CoV-2 variants, the Touret F and Ulrich L groups confirmed that the Alpha variant exhibits better replication fitness than the original viral strains in human reconstituted bronchial epithelium and animal models, respectively(11, 12). In addition, Liu Y and colleagues revealed that the Delta variant efficiently outcompetes the Alpha variant during coinfection in human lung epithelial cells and primary human airway tissues(13). These results align with the actual situation of the SARS-CoV-2 pandemic, in which the Alpha variant outcompeted WT SARS-CoV-2, and the Delta variant replaced the Alpha variant. However, Yuan S and colleagues recently demonstrated that the Omicron variant has no fitness advantage over Delta, which is inconsistent with that the former variant outcompeted the latter in the real world(14). These findings illustrate that the effects of the viral fitness for the competition among SARS-CoV-2 variants is not always the same. Immune pressure represents another driving factor that promotes the generation of novel variants of some pathogenic viruses(15-18). Recently, Ravindra K Gupta confirmed that convalescent plasma treatment for immune-suppressed individuals led to the emergence of SARS-CoV-2 variants with reduced susceptibility to neutralizing antibodies(19). However, whether immune selection pressure contributes to the emergence and displacement of SARS-CoV-2 epidemic variants lacks direct evidence. The competitive fitness among SARS-CoV-2 and its variants remains undetermined, especially under different statuses of immune pressure.

Homologous recombination has been documented in many human coronaviruses, including SARS-CoV, MERS-CoV, OC43, NL63, and HKU1(20-22), and in feline and canine coronaviruses(23, 24). Bioinformatic analysis has suggested frequent homologous recombination among SARS-CoV-2 variants, including Alpha and Delta(25), Delta and Omicron(26-29), and even Omicron sublineages(30, 31). Notably, some of the recombinants, such as recombinant XBB, have shown enhanced immune escape capability in comparison with their parental lineages(32, 33) and have even gained dominance in Singapore, India and other regions of Asia(34), indicating that SARS-CoV-2 recombination has great potential to produce new waves of infections and poses a threat to human populations. However, all previous findings are based on bioinformatics analyses, and no experimental evidence of recombination of SARS-CoV-2 has been documented.

In the present study, we characterized the competitive fitness of SARS-CoV-2 and its subsequent variants in two susceptible cell lines with or without immune pressure by using single nucleotide analysis combined with next-generation sequencing (NGS). More importantly, single clone purification of the passaged viruses confirmed the occurrence of homologous recombination among SARS-CoV-2 and its variants.

## Results

### Distinct competition patterns of SARS-CoV-2 and its progeny variants in Vero E6 and Caco-2 cells

To characterize the competitive fitness of SARS-CoV-2 and its progeny variants, direct competition experiments were performed by using the WT SARS-CoV-2, Alpha, Beta and Delta variants. Two SARS-CoV-2-susceptible cell lines, human colorectal carcinoma Caco-2 and African green monkey kidney Vero E6, were used in the present study. Briefly, a mixture of the four viruses at a 50% tissue culture infectious dose (TCID_50_) ratio of 1:1:1:1 was inoculated into each cell line, and the supernatants were collected and passaged for an additional 4 passages (six replicates were included at each passage). The original viral mixture was named P0, and the passaged viruses were named P1 to P5 (Fig 1a). The samples from each passage were quantified by RT‒qPCR, and no significant difference in total viral RNA loads was detected among different passages in both Vero E6 and Caco-2 cells (Extended Data Fig 1). To identify the ratio of each virus in the samples, we first selected a list of alleles specific to each of the four viruses by sequencing and alignment (Fig 1b) and then performed NGS and single nucleotide analysis to assess the frequency of each virus-specific alternate allele (Fig 1b) as previously described(35-37). In Caco-2 cells, for all six replicates, the frequency of the beta variant alleles rapidly became predominant at P1, while the frequency of the other three virus-specific alleles sharply decreased to almost undetectable levels by P4, indicating that the beta variant possesses stronger replication fitness and outcompetes the other three strains in Caco-2 cells (Fig 1c). However, in Vero E6 cells, the frequency of Beta and Delta alleles readily decreased and became undetectable by P3 in all six replicates, while the WT and Alpha alleles became predominant, except for two WT-specific alleles (Fig 1d), indicating that the WT and Alpha viruses possess stronger replication fitness than the other two variants in Vero E6 cells. The Beta variant showed enhanced fitness in human Caco-2 cells but reduced fitness in Vero E6 cells, while the WT and Alpha viruses showed opposite patterns in the two cell lines, suggesting that the competitive fitness of different SARS-CoV-2 variants varied in a cell-specific manner.

**Fig. 1.**
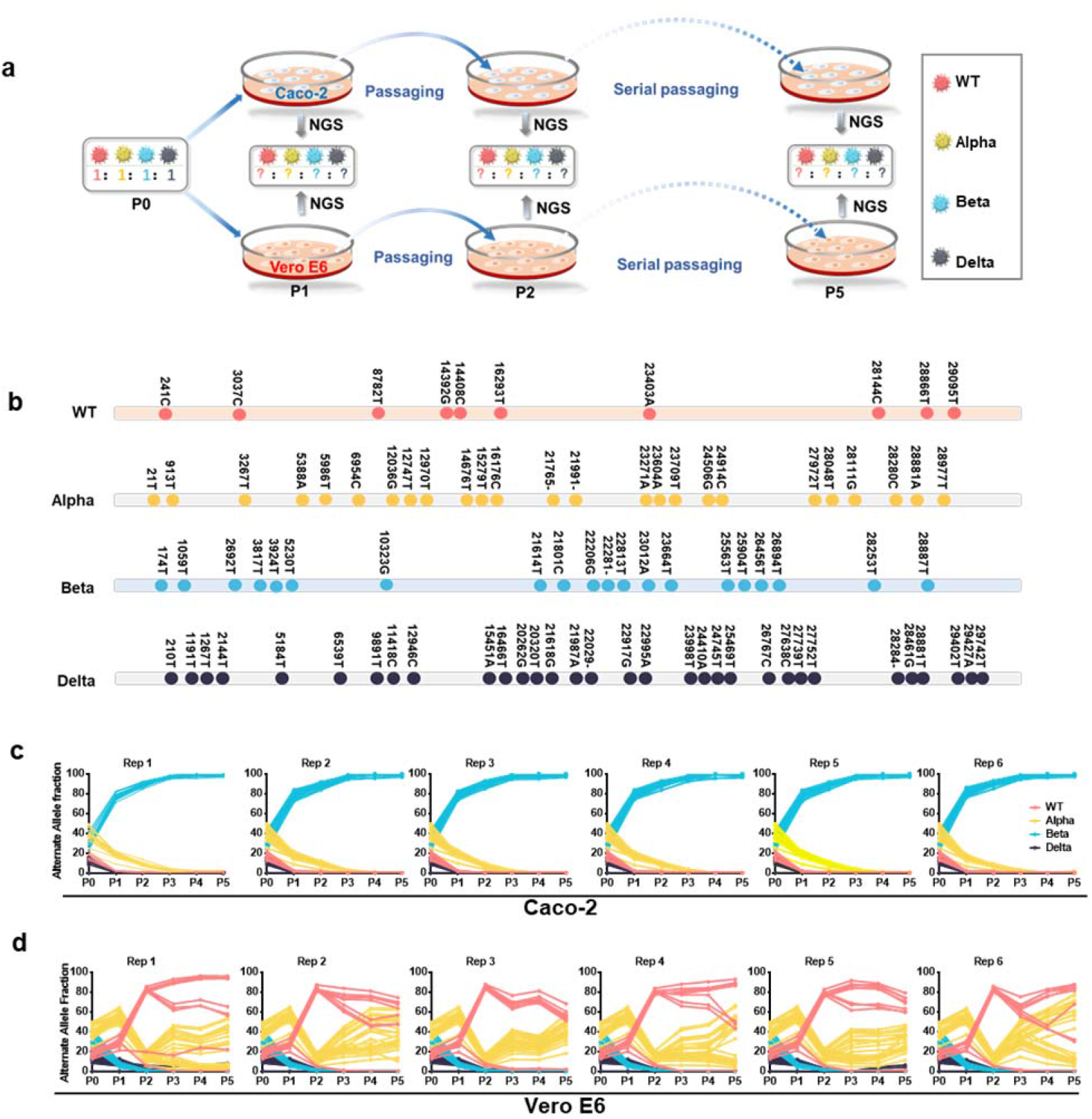
Competitive fitness of multiple SARS-CoV-2 strains in Vero E6 and Caco-2 cells. (a) Schematic diagram of the coinfection and passage history of SARS-CoV-2. The wild-type (WT) SARS-CoV-2, Alpha, Beta and Delta variants were mixed at equal TCID_50_ proportions (regarded as P0) to coinfect and serially passage Vero E6 or Caco-2 cells for five rounds. The samples collected from the passages were referred to as P1-P5. Six biological replicates were included for each passage, and the samples obtained from each replicate during the passage were analyzed by next-generation sequencing (NGS) to determine the frequency of each virus. (b) Schematic of specific alleles selected for each virus confirmed by sequencing and alignment. (c-d) The dynamic frequency of specific alleles for each of the four viruses at each passage in Caco-2 (c) and Vero E6 cells (d). Each color represents a SARS-CoV-2 strain, and each line represents a specific allele.

### Immune pressure reshaped the competition patterns of SARS-CoV-2 and its progeny variants

To further determine whether immune pressure affects the competitive fitness of SARS-CoV-2, the aforementioned direct competition assays were performed in the presence of high or low levels of neutralizing antibodies (Fig 2a). Three biological replicates were included for each passage. Under a relatively lower level of immune pressure, the WT and Alpha viruses rapidly outcompeted the Beta and Delta viruses, and the overall competition patterns and changing trends were similar to the original findings without antibody treatment (Fig 2b). However, under a relatively higher level of immune pressure, the Beta virus rapidly became predominant, and the other three viruses rapidly decreased to almost undetectable levels after P2 (Fig 2c). These results indicate that immune pressure obviously changes the original competition patterns of SARS-CoV-2 and its variants, and the Beta variant was the more potent variant under immune pressure.

**Fig. 2.**
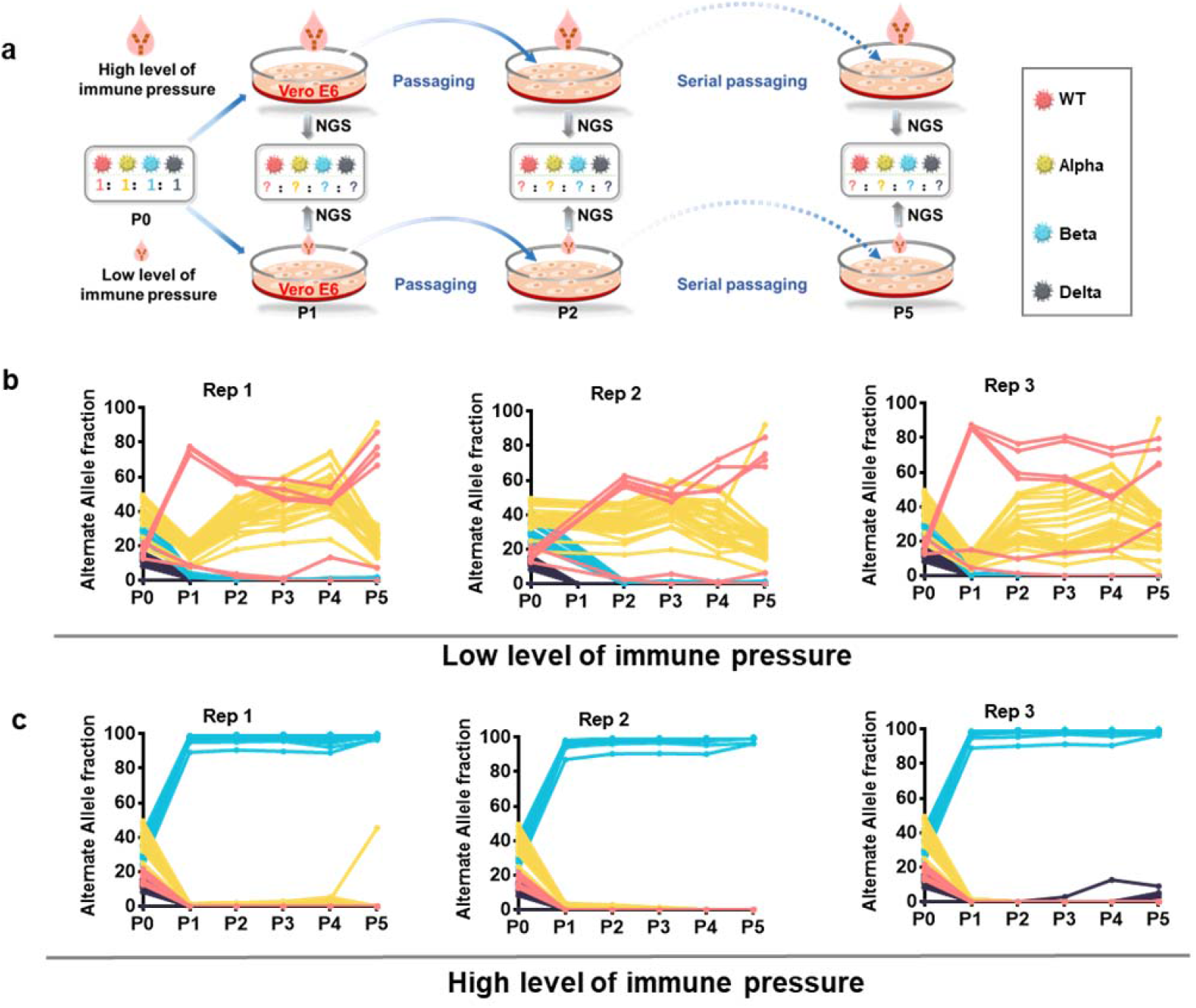
The competition of multiple SARS-CoV-2 viruses under different levels of immune pressure. (a) Schematic diagram of the coinfection and passage history of SARS-CoV-2 viruses under two levels of immune pressure. WT, Alpha, Beta and Delta were mixed at equal TCID_50_ proportions for coinfection and serial passage in Vero E6 cells for five rounds. Low and high immune pressure levels were maintained by adding 1 or 10 μl of vaccinated sera (NT_50_=1/512) during passage. Three biological replicates were included for each passage, and the samples obtained from each replicate were analyzed by NGS to determine the frequency of each virus. (b-c) The dynamic frequency of specific alleles for each of the four strains at each passage under low (b) and high (c) levels of immune pressure. Each color represents a SARS-CoV-2 virus, and each line represents a specific allele.

### Coinfection with SARS-CoV-2 and its variants led to homologous recombination

As shown in Fig. 1d and Fig. 2b, the WT and Alpha viruses coexisted throughout the passaging history in Vero E6 cells with or without neutralizing antibodies, which provided opportunities for homologous recombination. To further verify whether sustained coinfection could lead to homologous recombination, viral plaque purification was performed for the final P5 viral stocks (Fig 3a). In total, 28 and 10 plaques were cloned and amplified from the P5 stocks under no and relatively lower levels of immune pressure, respectively. The amplified viruses were then subjected to NGS and single nucleotide analysis (Fig 3a). Among the former 28 clone-purified SARS-CoV-2, 20 clones were identified as the WT virus, 2 clones were identified as the Alpha variant, and the remaining 6 clones (named Clone X1 to X6) were not completely identical to any of the four loading viruses (Fig 3b). Among the latter 10 clone-purified SARS-CoV-2, 7 clones’ specific alleles were identical to WT, and 2 clones’ specific alleles were identical to the Beta variant, but the specific alleles of the remaining 1 clone (named Clone X7) were not completely identical to any of the four viruses (Fig 3c). For Clone X1, the alleles are Alpha-specific from the 5’ end of the genome to nucleotide position 24914, while the Alpha-specific alleles all disappeared after nucleotide position 27972. In contrast, the alleles in the rest of the genome of Clone X 1 became WT specific. Similarly, Clones-X2 and X3 carried both Alpha- and WT-specific alleles near the 5’ end and 3’ end of the genome, with breakpoints located in regions of nucleotide positions 24506-24914 and 3267-5388, respectively (Fig 3c). These characteristics indicate that clones X1-X3 are SARS-CoV-2 recombinants between Alpha and WT (Fig 3c). For Clone X4, the alleles were WT-specific from the 5’ end of the genome to nucleotide position 26456, while the alleles changed to Beta-specific from nucleotide position 26894 to the 3’ end of the viral genome, indicating that this clone is a recombinant between WT and Beta with a breakpoint located in the region between nucleotide positions 26456 and 26894 (Fig 3c). Regarding Clones X5 and X6, most of the alleles were WT specific, but the WT-specific alleles at nucleotide position 23403 disappeared, and two Delta-specific alleles were also found at nucleotide positions 23998 and 24410 of the genome (Fig 3c), indicating that these two clones are recombinants between WT and Delta with two breakpoints located in regions of nucleotide positions 23604-23998 and 24410-24745. For Clone X7, the alleles were WT-specific from the 5’ end of the genome to nucleotide position 28280, while the alleles changed to Alpha-specific from 28866 to the 3’ end of the viral genome, indicating that this clone is a recombinant between WT and Beta with a breakpoint located in the region between nucleotide positions 28280 and 28866 (Fig 3c). These results clearly demonstrate that different patterns of SARS-CoV-2 recombinants, including Alpha-WT, WT-Beta and WT-Delta, were generated during the coinfection and serial passaging history. Notably, among these 7 clones, 6 clones’ genome breakpoints were located between nucleotide positions 22995 and 28866, which is in line with SARS-CoV-2 recombination hotspots identified by previous case reports or bioinformatic analyses.

**Fig. 3.**
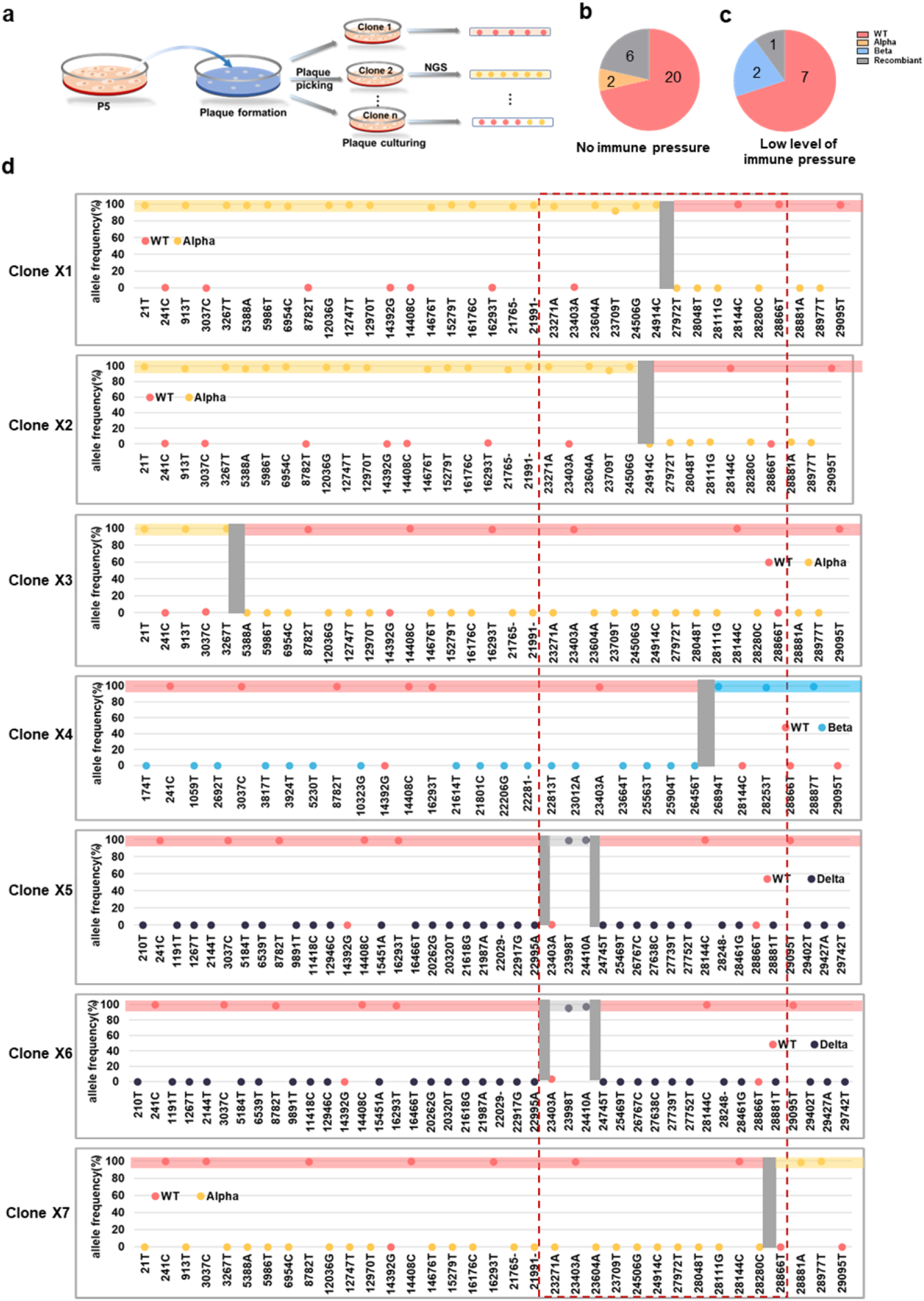
Coinfection with multiple SARS-CoV-2 viruses led to recombination. (a) Schematic diagram of the recombination analysis for P5 samples by plaque isolation and sequencing. (b-c) The overall genome characteristics of the plaques isolated and cloned from P5 samples obtained from coinfection and serial passage under no (b) or low levels of immune pressure (c). (d) Graph representing the alternate allele frequency of each specific allele of the 7 clones with recombination characteristics. Alleles specific to WT, Alpha, Beta and Delta are represented in pink, yellow, blue and black, respectively. The gray box represents the region where the breakpoint occurred in the genome of each clone. The red box represents the region where the breakpoints preferentially occur (22995-28866).

## Discussion

In this study, we revealed that not only viral fitness but also immune pressure and host species variation drove the competition of multiple SARS-CoV-2 variants, providing insight into the competition and replacement mechanism of epidemic SARS-CoV-2 variants in the real world.

First, we provide direct evidence that the Beta variant has competitive fitness over WT SARS-CoV-2 in human cells (Fig. 1c). Although no advantage was observed in Vero E6 cells, the Beta variant outcompeted the other three variants within two passages in human Caco-2 cells. This result partially explains the scenario in which WT SARS-CoV-2 was outcompeted in humans in South Africa and several other countries during early 2021(3). However, unexpectedly, the Delta variant was outcompeted by the Beta variant in Caco-2 cells during coinfection and serial passaging, which is contrary to the scenario in which the Delta variant replaced the Beta variant worldwide. This result is consistent with the results of a recent study showing that the Omicron variant did not show higher viral fitness than the Delta variant, but the former indeed displaced the latter in the real world(14). These results indicate that the competitive fitness of some variants may be influenced by other factors in addition to replication fitness.

Additionally, the distinct competition outcomes of SARS-CoV-2 variants in different cells deserve specific attention. Diverse competition outcomes of SARS-CoV-2 viruses in different experimental animals have been revealed recently (12). As multiple SARS-CoV-2 variants have infected domesticated and wild animals and even established sustained transmission (38-40), the competition of multiple SARS-CoV-2 variants in these new host species will be inevitable, and the predominant variants that finally appear in these species may also be different. As the cross-species transmission of SARS-CoV-2, including human-to-animal-to-human transmission(40), has been reported, different predominant SARS-CoV-2 variants simultaneously circulating in multiple species may cause new risks, such as coinfection and recombination of multiple variants. Regardless, along with the continual circulation and adaptation of SARS-CoV-2, the competitive fitness of these new variants in diverse hosts deserves further investigation in the future.

More importantly, our study showed that the competitive fitness of the Beta variant increased under immune pressure. As shown in Fig. 2c, a high level of neutralizing antibodies enabled the disadvantaged Beta variant to rapidly outcompete other viruses in Vero E6 cells. Previously, it was demonstrated that the Beta variant evolved a stronger immune escape capability against infected and vaccinated sera than the other three viruses(41). Thus, the dramatic change in the competition outcomes of the four SARS-CoV-2 viruses after immune pressure was added is probably attributed to the different immune escape capabilities obtained by these viruses. This phenomenon also indicates the key role of the immune barrier of humans in the emergence and displacement of predominant variants.

Most significantly, our plaque purification experiments provided direct evidence supporting homologous recombination among different SARS-CoV-2 variants during coinfection. All previous findings regarding homologous recombination in SARS-CoV-2 are based on sequence alignment and bioinformatics studies(42-45). In our study, by single plaque purification, a direct and classical procedure for recombinant analysis, 6 of 28 clones (∼21%) were identified as recombinants from coinfected parental viruses, providing experimental evidence of SARS-CoV-2 recombination for the first time. Meanwhile, the unexpectedly high rate of recombinants we observed in vitro also indicates that the recombination of SARS-CoV-2 viruses is prone to occur during coinfection. However, understanding the real scenario of recombination in vivo requires further investigation.

SARS-CoV-2 recombination has caused great concern with the emergence and transmission of XBB. XBB is a recombinant between the variant BJ.1 and BM.1.1.1 with the breakpoint located in the S gene (22897-22941). By recombination, XBB gained more mutations in the S gene, including those able to enhance immune evasion (Y144del and F486S) and infectivity (V83A and N460K), and showed higher immune evasion than BA.2, BA.4/5 and even BQ.1/1.1(32, 46, 47). Currently, this recombinant is rapidly spreading worldwide and has already become the predominant variant in several Asian countries and America by outcompeting BA.2, BA.5 or even BQ.1, showing higher competitive fitness than their parental strains and even other Omicron sublineages (34). Thus, investigating the process and mechanism of SARS-CoV-2 recombination is important and urgent. Of note, the breakpoints of 6 of the 7 recombinants we obtained in this study are between nucleotide positions 22995 and 28866, consistent with the recombination hotspots identified by case reports or inferred by different phylogenomic methods(42-45). Notably, the breakpoints of Clones X2 and X5/6 are located in the S gene (24506-24914, 23604-23998 and 24410-24745) (Fig 3d and Extended Data Fig 2), which are similar to the recombination breakpoints of XBB (Extended Data Fig 2). Thus, the strategy of coinfection and serial passage with different vrial strains in vitro adopted in our study mimicked SARS-CoV-2 recombination in humans, which will be a very helpful model for investigating the recombination mechanism.

In conclusion, our study profiled the variable competitive fitness of SARS-CoV-2 under different conditions and deepened our understanding of the complexity of competition and replacement of SARS-CoV-2 epidemic variants in the real world. Meanwhile, we successfully built an in vitro model of reproducing the recombination of SARS-CoV-2 under experimental conditions for the first time, which will be helpful for further investigating SARS-CoV-2 recombination mechanism.

**Extended Data Fig. 1.**
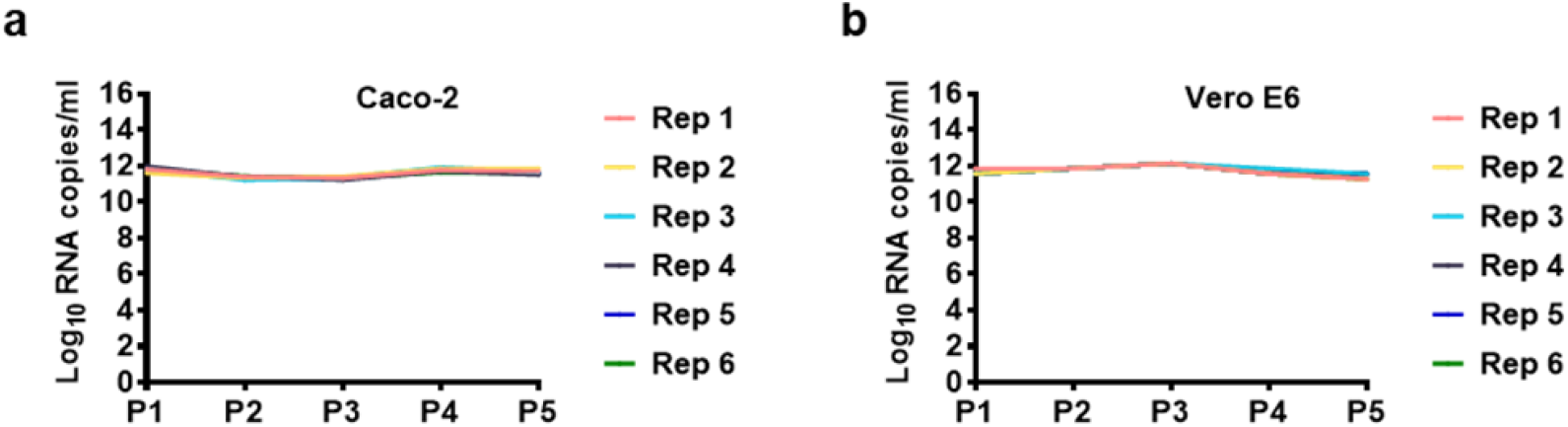
RNA copies of SARS-CoV-2 during serial passage in Vero E6 and Caco-2 cells. One hundred microliters of the viral stock obtained from each passage was adopted for viral RNA extraction and quantified by RT‒qPCR. (a) The viral RNA copies in samples obtained from passages of Caco-2 cells. (b) The viral RNA copies in samples obtained from passages of Vero E6 cells.

**Extended Data Fig. S2.**
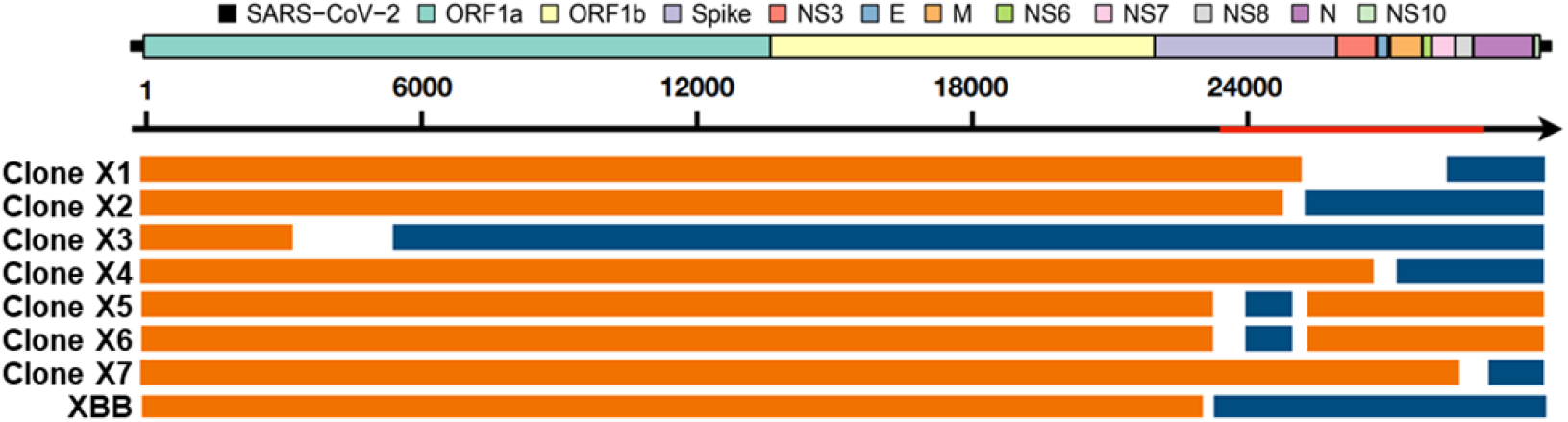
Schematic diagram of the genomes of recombination clones and XBB.

## Methods

### Cell lines and viruses

The SARS-CoV-2 strain IME-BJ05 (accession no. GWHACAX01000000) used as the wild type SARS-CoV-2 (WT) in this study was originally isolated from a COVID-19 patient in China early 2020(48). The Alpha variant (C-Tan-BJ202101) was from Chinese Center for Disease Control and Prevention(49). The Beta variant GDPCC (CSTR: 16698.06. NPRC 2.062100001) was from the National Pathogen Resource Center (NPRC), China(50). The Delta variant (CCPM-B-V-049-2105-8) was from Chinese Academy of Medical Sciences(49). SARS-CoV-2 was amplified and titrated by Median Tissue Culture Infectious Dose (TCID_50_) in Vero E6 cells. Vero E6 cells (CCL-81) and Caco-2 cells (HTB-37) were from ATCC, and maintained in Dulbecco’s minimal essential medium (DMEM, Thermo Fisher Scientific) supplemented with 10% fetal bovine serum (FBS, Thermo Fisher Scientific) and penicillin (100 U/ml)-streptomycin (100 μg/ml) (Thermo Fisher Scientific). All experiments involving infectious SARS-CoV-2 were performed in biosafety level 3 (BSL-3) containment laboratory in Beijing Institute of Microbiology and Epidemiology, AMMS.

### Coinfection and serial passage of SARS-CoV-2 viruses in Vero E6 cells or Caco-2 cells

Confluent monolayers of each cell line were prepared for infection by seeding six-well plates with 5 × 10^5^ cells/well in 2 ml of media and incubating at 37 °C for 12 hours. WT, Alpha, Beta and Delta viral stocks were mixed at a TCID_50_ ratio of 1:1:1:1 and diluted to 5 × 10^5^ TCID_50_ / ml. 100 μl of the mixture was inoculated into each well and then incubated at 37 °C. The medium from each well was harvested at 72 hours post infection, and 100 μl of which was used to serially passage in relative cells for another four rounds and the rest medium obtained at each passage was stored at −80°C for quantification and sequencing.

### Coinfection and serial passage of SARS-CoV-2 viruses in Vero E6 cells or Caco-2 cells under immune pressure

Ten serum samples from vaccinees received two doses of an mRNA vaccine ARCoV, which encodes the SARS-CoV-2 spike protein receptor-binding domain (RBD)(51) were collected and neutralizing antibody titers were tittered by a standard plaque reduction neutralization test (PRNT) as described previously(52). The neutralization titers of serum samples were all diluted to ∼1/512 and then mixed at equal volume. Coinfection and serial passage of SARS-CoV-2 viruses were performed as above, and each well was further added with 1 or 10 μl prepared serum mixture (which were regarded as low or high dose of neutralizing antibodies, respectively) at each passage to form a relatively lower or higher level of immune pressure, respectively.

### RNA Extraction, quantification and Sequencing

The RNA of all cell culture supernatants was extracted using High Pure Viral RNA kit (Roche, REF.11858882001). Viral RNA quantification in each sample was performed by quantitative reverse transcription PCR (RT-qPCR) targeting the S gene of SARS-CoV-2. RT-qPCR was performed using One Step PrimeScript RT-PCR Kit (Takara, Japan) with the following primers and probes: CoV-F3 (5’-TCCTGGTGATTCTTCTTCAGGT-3’); CoV-R3 (5’-TCTGAGAGAGGGTCAAGTGC-3’); and CoV-P3 (5’-AGCTGCAGCACCAGCTGTCCA-3’). The sequencing libraries were constructed using the MGIEasy RNA Library Prep Set V3.0 (MGI, No. 1000006383) kit for paired-end sequencing (2×150 bp reads). The library was sequenced using MGISEQ2000 sequencing platform.

### High Throughput Sequencing Data Analysis

The sequencing data filtering was performed using fastp v0.21.0(53). Then we used CLC Genomics Workbench v21.0.4 to do reads mapping and single nucleotide variants (SNV) calling. The fitered sequences were mapping to SARS-CoV-2 reference sequence (Wuhan-Hu-1, GenBank accession number NC_045512.2). The mapped length fraction threshold was set to 90%. The similarity threshold was set to 90%. The single nucleotide polymorphism (SNP) was called according to mapping results. All site which coverage lower than 3, mutation count lower than 2, or mutation frequency lower than 1% were excluded in SNP calling. The genome sequences of clone X1-clone X7 have been submitted to NCBI databases and the accession numbers were OQ231280, OQ231281, OQ231502, OQ217008, OQ231282, OQ231283 and OQ231505, respectively.

## Acknowledgements

This study was supported by the National Key Research and Development Projects of the Ministry of Science and Technology of China (2021YFC2301300, 2021YFC2301302) and National Natural Science Foundation of China (NSFC) (No.32130005). CFQ was supported by the National Science Fund for Distinguished Young Scholars (81925025), the Innovative Research Group (81621005) from the NSFC, and the Innovation Fund for Medical Sciences (2019-I2M-5-049) from the Chinese Academy of Medical Sciences. HF was supported by the Beijing Nova Project (Z201100006820029)

## Author contributions

Cheng-Feng Qin and Qi Chen conceived and designed the project. Qi Chen and Yong-Qiang Deng performed the coinfection and serial passage experiments. Si Qin, Hang Fan, Hang-Yu Zhou, Pan-Deng Shi, Hui Zhao, Xiao-Feng Li, Xing-Yao Huang, Ya-Rong Wu, Yan Guo, Guang-Qian Pei, Yun-Fei Wang performed the NGS experiments and analyzed the data. Cheng-Feng Qin, Qi Chen, Si Qin, Hang Fan, Hang-Yu Zhou, Zong-Min Du and Yu-Jun Cui wrote and finalized the manuscript. All authors read and approved the manuscript.

## Competing interests

The authors declare no competing interests.

